# Ablation of LRP6 in alpha-smooth muscle actin-expressing cells abrogates lung inflammation and fibrosis upon bleomycin-induced lung injury

**DOI:** 10.1101/2024.09.05.611327

**Authors:** Eun-Ah Sung, Mikhail G. Dozmorov, SuJeong Song, Theingi Aung, Min Hee Park, Patricia J. Sime, Wook-Jin Chae

## Abstract

Low-density lipoprotein receptor-related protein 6 (LRP6) is a receptor for Wnt ligands. Tissue fibrosis is a progressive pathological process with excessive extracellular matrix proteins (ECM) deposition. Myofibroblasts, identified by alpha-smooth muscle actin (αSMA) expression, play an important role in tissue fibrosis by producing ECM production. Here we found that Wnt antagonist Dickkopf1 (DKK1) induced gene expressions associated with inflammation and fibrosis in lung fibroblasts. We demonstrated that genetic deletion of LRP6 in αSMA-expressing cells using *Acta2*-cre *Lrp6*^fl/fl^ (*Lrp6*^AKO^) mice abrogated bleomycin (BLM)-induced lung inflammation and fibrosis phenotype, suggesting an important role of LRP6 in modulating inflammation and fibrotic processes in the lung. Our results highlight the crucial role of LRP6 in fibroblasts in regulating inflammation and fibrosis upon BLM-induced lung injury.

## Introduction

Environmental challenges can cause tissue damage and trigger inflammatory immune response [1]. These immune responses are closely integrated with tissue repair and remodeling, facilitating the preservation of tissue homeostasis [2]. Dysregulation of these processes can result in chronic inflammation and ultimately lead to tissue fibrosis.

The Wnt signaling pathway is crucial for cell differentiation, proliferation, self-renewal and immune cell regulation for tissue repair and regeneration. Low-density lipoprotein receptor-related protein 6 (LRP6) acts as a co-receptor for Wnt ligands that are essential for activating the canonical Wnt signaling pathway [3]. LRP6 is a transmembrane protein and belongs to low-density lipoprotein receptor (LDLR) family. In the presence of the Wnt agonist Wnt3a, Dickkopf1 (DKK1) binds competitively to LRP6 with higher affinity, inhibiting the canonical Wnt signaling pathway [4,5]. Given that dysregulation of canonical Wnt signaling causes multiple inflammatory diseases, it is increasingly clear that Wnt pathway mediators and ligands play important immune-modulatory roles [3,6,7].

Pulmonary fibrosis (PF) is a progressive disease marked by fibroproliferation due to imbalanced injury-inflammation-repair processes. Bleomycin (BLM)-induced lung injury model is a widely used mouse model for studying PF. BLM, a profibrotic agent, can cause PF when administered intravenously for lymphoma treatment [8]. BLM-mediated lung injury triggers lung inflammation and fibrosis, mimicking the accelerated acute phase of PF observed in humans [9–11]. Elevated levels of DKK1 have been shown in both human PF patients and BLM-induced lung fibrosis in mice [12–14]. Reduced expression of DKK1 in *doubleridge* mice, DKK1 neutralization and thrombocyte-specific deletion of DKK1 protected the mice from BLM-induced lung injury, reducing inflammation and fibrosis phenotypes [14,15]. PF is caused by the deposition of extracellular matrix (ECM) proteins [16]. While LRP6 is a key receptor for DKK1, its role in lung fibroblasts within PF pathology remains unclear.

PF is characterized by excessive proliferation of myofibroblasts, driven by increased profibrotic cytokines, and accumulation of ECM proteins, such as collagen [17]. Myofibroblasts, identified by their expression of alpha-smooth muscle actin (αSMA), play a crucial role in producing ECM proteins that provide the mechanical properties necessary for the organ’s function [18]. In BLM-induced lung injury, αSMA-expressing cells, such as myofibroblasts, accumulate in fibrotic areas and contribute to collagen deposition [19,20]. Aberrant activation or differentiation of fibroblasts dysregulates homeostatic crosstalk with other immune cell types, such as macrophages, for tissue repair, leading to fibrosis [21–23]. Inhibition of myofibroblast differentiation has been shown to reduce or abrogate PF progression [24–26].

In this study, we found that DKK1 induced gene expression profiles for inflammation and tissue repair in lung fibroblasts. We addressed the role of DKK1 receptor LRP6 in αSMA-expressing cells using *Acta2*-cre *Lrp6*^fl/fl^ mice in the BLM-induced lung injury model. Our results showed that LRP6 ablation in αSMA-expressing cells markedly reduced BLM-induced lung inflammation and fibrosis. We identified the importance of LRP6 in αSMA-expressing cells as a critical mediator in regulating inflammation and tissue repair upon BLM-induced lung injury.

## Materials and Methods

### Mouse lung fibroblast isolation and purification

Lungs were perfused, then chopped into small pieces and digested in a collagenase solution (Collagenase type IV (1200 U/ml), 2% FBS and 2% Pen/Strep in sterile PBS). After a 30-minute incubation at 37 °C, the homogenates were centrifuged at 1000 x g for 5 minutes at room temperature, then washed in a washing solution (2% FBS and 2% Pen/Strep in sterile PBS). The homogenates were resuspended in 0.05% Trypsin-EDTA and incubated at 37 °C for 20 minutes. After centrifugation, cells were suspended in fibroblast culture medium (DMEM supplemented with 10% FBS, 2% Pen/Strep and 1% GlutaMax) and seeded into 6-well plates. After seven days, the cells were detached and purified. For the purification step, cells were incubated with biotinylated antibodies against EpCAM (118203, BioLegend), CD31 (102503, BioLegend), and CD45 (13-0451-82, eBioscience), followed by the addition of MojoSort Streptavidin Nanobeads (480015, BioLegend). Cells were separated using a MojoSort Magent (480019, BioLegend). Sorted cells were then collected for further analysis.

### Bulk RNA seq

Total RNA was extracted from lung fibroblasts using TRIzol (15596026, Thermo Fisher Scientific), followed by purification with the Rneasy Plus Mini Kit (74136, QIAGEN), according to the manufacturer’s instructions. RNA-seq data was processed using Nextflow rnaseq pipeline (v.3.14) which performs QC, adapter trimming, alignment to reference genome GRCm38 using STAR [27] and transcript quantification using RSEM [28]. Differential gene expression (DEG) analysis was conducted using Bioconductor package edgeR v4.2.1, based on normalized and filtered gene counts [29]. Genes exhibiting statistically significant changes (adjusted p-value < 0.05) were further analyzed using downstream bioinformatics approaches. The DEGs are listed in **Table S1**.

### Animal experiments

*Acta2*-cre mice (#032758) and *Lrp6* flox mice (#026267) were purchased from the Jackson Laboratory and have been bred in our mouse facility. All mouse protocols were approved by the VCU Animal Care and Use Committee (IACUC) and followed the guidelines of the Association for Assessment and Accreditation of Laboratory Animal Care International (AAALAC).

For the bleomycin-induced lung injury model, Bleomycin Sulfate (BLM; S121415, Selleckchem) (4 U/kg) was dissolved in PBS and administered via the oropharyngeal route. Control mice were treated with PBS. Eight-week-old *Acta2*-cre *Lrp6*^fl/fl^ and *Lrp6*^fl/fl^ mice were challenged with BLM on day 0, and the lungs were harvested on day 14.

### Hemavet analysis

Peripheral blood was collected from *Lrp6*^fl/fl^ and *Acta2*-cre *Lrp6*^fl/fl^ mice using the retro-orbital sinus sampling method. The Hemavet 950FS (Drew Scientific) was used for the analysis.

### Immunohistochemistry (IHC)

Lungs were fixed with 10% formalin (23-245685, Fisher Scientific) and embedded in paraffin. Lung tissue sections (5 μm thickness) were used. H&E staining and Masson’s Trichrome staining were performed as described previously [14]. For αSMA IHC, endogenous peroxidases were blocked using 3% H_2_O_2_. The sections were then incubated with a primary antibody against αSMA (1:200; PA5-16697, Thermo Fisher Scientific). Following this, the sections were stained with 5 μg/ml of an HRP-conjugated secondary antibody. The staining was developed using a DAB substrate kit (SK-4100, Vector Laboratories), and the nuclei were counterstained with Hematoxylin. Images were acquired using Vectra Polaris (Akoya Biosciences).

### Quantification of IHC images

The acquired images were saved in TIFF format using Phenochart (Akoya Biosciences). Image processing was performed using the “Color Deconvolution” plugin in Fiji software (H DAB for αSMA IHC). Images were deconvoluted into their respective color components. The brown component of H DAB was measured using the “Threshold” tool. The Threshold value for image analysis was adjusted and consistently applied to all processed images. The selected areas were analyzed using the “Measure” tool.

### Flow cytometry

Single-cell lung homogenates were prepared using collagenase (280 U/ml, Collagenase type IV (LS004188, Worthington Biochemicals), 2 % (v/v) FBS and DNase I (40 μg/ml, 10104159001, Roche) in PBS using the gentleMACS Octo Dissociator (Miltenyi Biotec). After blocking Fc Receptor, the cells were stained with Zombie Aqua Fixable Viability Kit (423102, BioLegend) and then with fluorescent-conjugated antibodies against various cell surface antigens. Flow cytometry was performed using FACSymphony A1 (BD Biosciences), and the data were analyzed with FlowJo software (version 10.7, Tree Star). Antibody information is shown in **Table S2**.

### Hydroxyproline Quantitation Assay

Lungs were harvested and processed to quantify hydroxyproline using a hydroxyproline assay kit (MAK008, Sigma-Aldrich). Following the manufacturer’s protocol, hydroxyproline concentration was measured using a colorimetric assay at 560 nm.

### Western blot

Lung homogenate lysates were separated by SDS-PAGE and transferred onto a polyvinylidene difluoride membrane. The membrane was blocked and then immunoblotted with primary antibodies against αSMA (1:1500, PA5-16697, Invitrogen), DKK1 (1:1500, AF1096, R&D Systems), IL-11 (1:1500, PA5-95982, Invitrogen) and Gapdh (1:3000, 5174S, Cell signaling Technology). The membrane was incubated with HRP-conjugated secondary antibodies for 1 hour at room temperature. The signal was developed using the WesternBright^TM^ Sirius kit (K-12043-D10, Advansta). Bands were detected and quantified using the Chemidoc and ImageJ software, and the results were normalized to Gapdh protein levels.

### Statistical Analysis

Statistically significant differences were determined using Student’s t-test or one-way ANOVA analysis, followed by Tukey’s post hoc test, performed with GraphPad Prism software (version 9.4.1, GraphPad Software Inc.). Data are presented as mean with standard deviation (SD). P values < 0.05 were considered statistically significant.

## Results

### DKK1 induces gene expression profiles for inflammation and tissue repair in lung fibroblasts

Given that DKK1 functions as a proinflammatory and profibrotic ligand promoting BLM-induced lung inflammation and fibrosis [14,15], we investigated its role in modulating lung fibroblast. To explore DKK1-mediated gene expressions, we used mouse lung fibroblasts and conducted bulk RNA seq. Kyoto Encyclopedia of Genes and Genomes (KEGG) pathway enrichment analyses revealed that DKK1-treated lung fibroblasts exhibited enrichment in essential components of the immune responses, including IL-17 signaling pathway, TNF signaling pathway, and cytokine-cytokine receptor interaction (**Fig. 1A**). Further analyses using Gene Ontology (GO) terms showed that DKK1 enhanced processes related to cytokine activity, regulation of inflammatory response, cytokine-mediated signaling pathway and regulation of defense response compared to untreated group (**Fig. 1B**). Bulk RNA seq analyses for differentially expressed genes (DEGs) revealed that DKK1 upregulated several genes that play key roles in inflammation and tissue fibrosis (**Fig. 1C, Fig. S1**).

**Figure 1.**
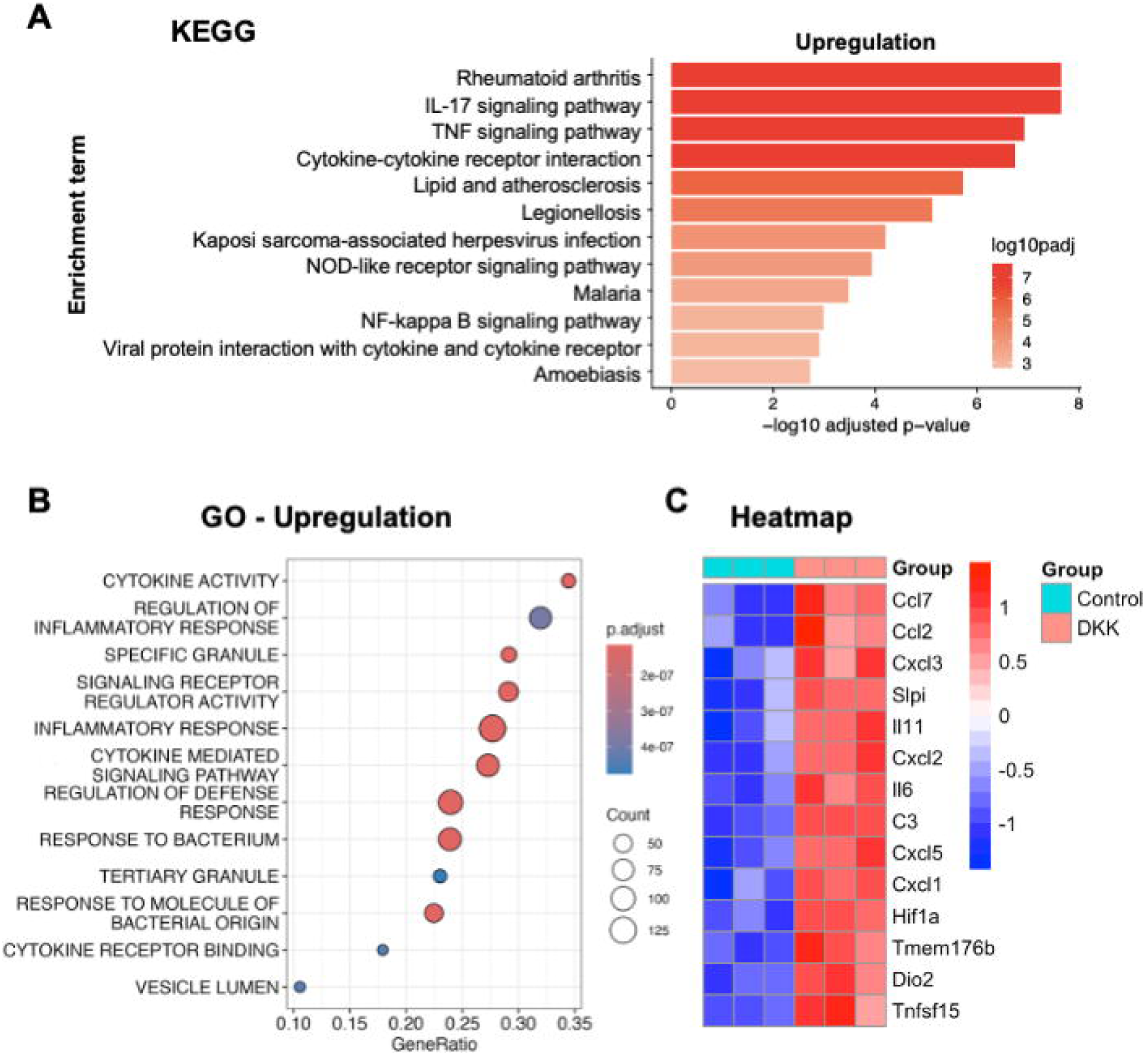
DKK1 induces gene expressions for inflammation and tissue repair in lung fibroblasts. (**A-C**) Lung fibroblasts were treated with DKK1 for 24 hours prior to bulk RNA seq. KEGG pathway enrichment analysis (**A**) and Gene Ontology (GO) enrichment analysis (**B**) were conducted to identify significantly upregulated pathways in DKK1-treated group compared to untreated group. DEG analysis between untreated and DKK1-treated group is visualized in the form of a heatmap (**C**).

DKK1 increased the expression of *C3* and *Dio2* genes in mouse lung fibroblasts (**Fig. 1C, Fig. S1**). Complement component 3 (C3) is upregulated in human PF lungs, and fibroblasts are a primary source of C3 in both human PF lungs and BLM-treated mouse lungs [30]. The activity and expression of deiodinase2 (DIO2) are higher in the lungs of PF patients compared to controls, correlating with disease severity [31].

DKK1 also increased the expression of *Il6* gene (**Fig. 1C, Fig. S1**). IL-6 levels are elevated in the lungs of PF patients and mouse models of PF [32,33]. IL-6-deficient mice exhibited attenuated lung inflammation and collagen deposition [33]. Additionally, DKK1 induced the expression of *Il11*, a member of the IL6 family of cytokines (**Fig. 1C, Fig. S1**). IL-11 is upregulated in human PF lungs and fibroblasts and activated lung fibroblasts with increased αSMA protein expression [34,35]. Recombinant mouse IL-11 injection or *Il11* overexpression in fibroblasts-or smooth muscle cell-specific manners induced lung fibrosis [35,36]. Fibroblast-specific deletion of *Il11ra1* or neutralization of IL-11 prevents BLM-induced lung fibrosis phenotypes [35,37,38]. DKK1 upregulated mRNA expression of chemokines, such as Cxcl1, Cxcl3, Cxcl5, Ccl2 and Ccl7 (**Fig. 1C, Fig. S1**). *Cxcl5*, which binds to Cxcr2, promoting the pathogenesis of PF [39].

DKK1 increased the mRNA expression of *Tmem176b*, also known as Lr8 (**Fig. 1C, Fig. S1**). Tmem176b expression is upregulated in human PF lungs and BLM-induced fibrotic mouse lungs [40]. LR8-expressing fibroblasts in tissues express αSMA [41]. Secretory leukocyte peptidase inhibitor (Slpi)-deficient mice showed reduced lung fibrosis upon BLM-induced lung injury [42]. mRNA expression of *Slpi* was increased by DKK1. *Tnfsf15* gene expression was increased by DKK1 (**Fig. 1C, Fig. S1**). Tnfsf15 gene encodes tumor necrosis factor-like cytokine 1A (TL1A). *Tnfsf15* mRNA expression was found to be elevated in isolated human lung fibroblasts stimulated with bronchoalveolar lavage fluid (BALF) from PF patients [43]. Additionally, the administration of recombinant TL1A increased fibrosis phenotypes, such as collagen deposition and αSMA^+^ cell population in the lung, and genetic deletion of its receptor DR3 reduced the fibrosis phenotypes upon BLM-induced lung injury [44].

Hypoxia has been shown to play an important role in lung fibrosis pathogenesis [45,46]. The hypoxia-inducible factor 1^α^ (Hif1α) pathway was upregulated in PF patient tissue samples and human lung fibroblasts exposed to a PF-conditioned matrix [46]. Fibroblast-specific *Hif1A* gene ablation reduced BLM-induced lung fibrosis [47]. DKK1 increased *Hif1a* mRNA expression (**Fig. 1C, Fig. S1**).

Collectively, our results indicated that DKK1 induces gene expression profiles associated with inflammation and tissue fibrosis in mouse lung fibroblasts.

### *Lrp6*^AKO^ mice showed normal hematological phenotypes

LRP6 is a key receptor for DKK1 and DKK1– and LRP6-deficiency have shown similar phenotypes in amphibian embryos [4,48,49]. Given the crucial role of αSMA-expressing cells in tissue repair through ECM production, we generated αSMA-expressing cell-specific *Lrp6*-deficient mice (*Acta2*-cre *Lrp6*^fl/fl^, hereafter referred to as *Lrp6*^AKO^). Isolated lung fibroblasts (CD45^-^EpCAM^-^CD31^-^ cells) from *Lrp6*^AKO^ and their littermate control *Lrp6*^fl/fl^ mice were used to examine LRP6 protein expression using flow cytometry (**Fig. S2**). LRP6 protein expression was analyzed in αSMA^+^ and αSMA^-^ cells. LRP6 protein expression was decreased in αSMA-expressing cells in *Lrp6*^AKO^ mouse lungs, but not in αSMA^-^ cells (**Fig. 2A** and **Fig. S2**).

**Figure 2.**
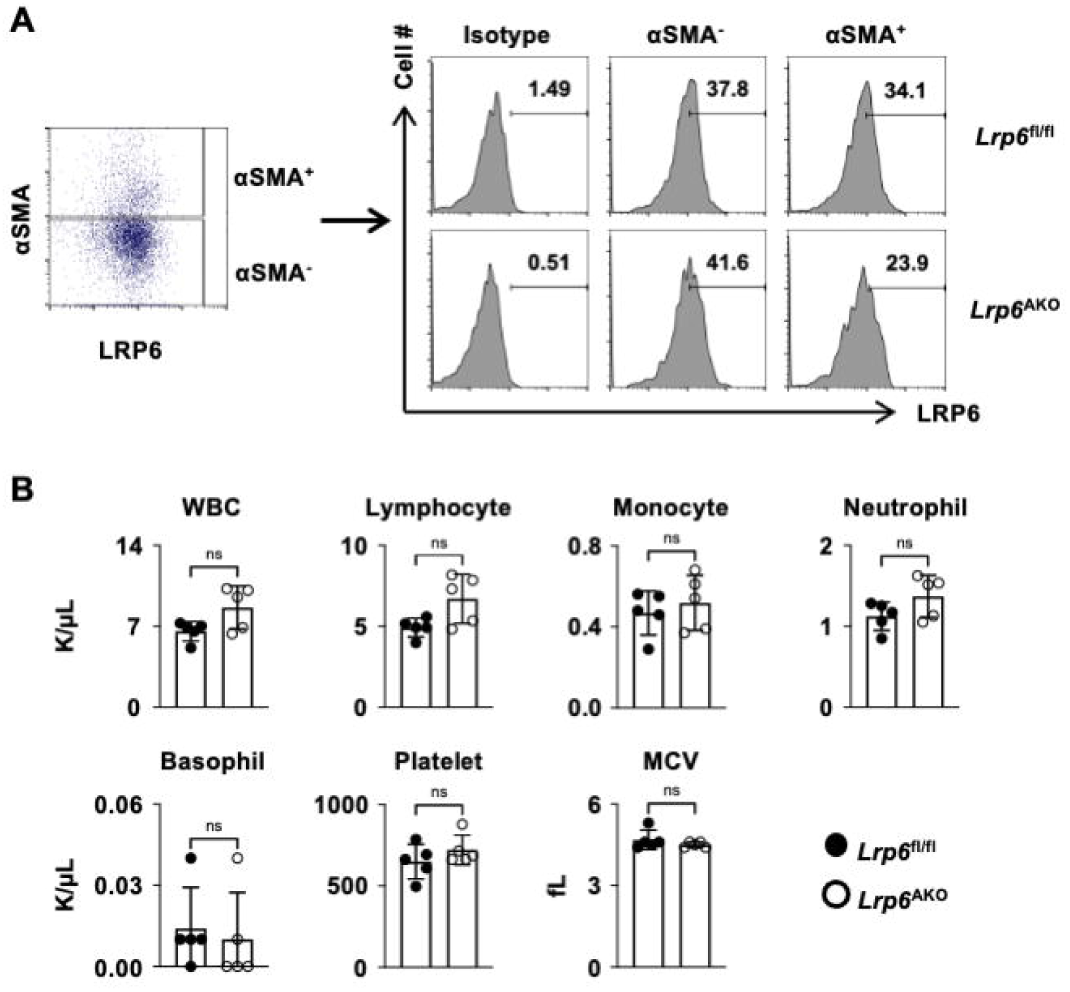
*Lrp6*^AKO^ mice showed normal hematological phenotypes. (**A**) LRP6 expressions in αSMA^+^ or αSMA^-^ populations were quantified by flow cytometry from the lungs of *Lrp6*^AKO^ and *Lrp6*^fl/fl^ mice. (B) Inflammatory cells in the peripheral blood from *Acta2*cre-*Lrp6*^fl/fl^ and *Lrp6*^fl/fl^ mice were analyzed by Hemavet. Shown are white blood cells (WBC), lymphocytes, monocytes, neutrophils, eosinophils, basophils, red blood cells (RBC), platelets and mean corpuscular volume (MCV) (n = 5). Data are presented as the mean ± SD. Student’s t-test. ns, not significant.

Next, we performed hematological analyses of *Lrp6*^AKO^ and *Lrp6*^fl/fl^ mice. We found minimal differences in peripheral blood hematological parameters between *Lrp6*^AKO^ and *Lrp6*^fl/fl^ mice, indicating normal hematological phenotypes in *Lrp6*^AKO^ mice (**Fig. 2B**). These results indicate that the ablation of LRP6 in αSMA-expressing cells did not alter baseline immune cell profiles in circulating blood under homeostatic condition.

### *Lrp6*^AKO^ mice showed markedly reduced collagen deposition in the BLM-induced lung injury model

To explore whether the ablation of LRP6 in αSMA-expressing cells reduced collagen deposition upon BLM-induced acute lung injury, *Lrp6*^AKO^ and their littermate control *Lrp6*^fl/fl^ mice were challenged with BLM on day 0. Two weeks later, lungs were analyzed using Masson’s Trichrome staining to detect collagen deposition. BLM-treated *Lrp6*^fl/fl^ mice showed a marked increase in collagen deposition compared to vehicle-treated *Lrp6*^fl/fl^ mice (**Fig. 3A, B**). Notably, *Lrp6*^AKO^ mice had significantly less collagen deposition upon BLM-induced lung injury compared to *Lrp6*^fl/fl^ mice (**Fig. 3A, B**). Consistent with these findings, hydroxyproline concentration in the lungs of *Lrp6*^AKO^ mice was decreased compared to *Lrp6*^fl/fl^ mice following the BLM challenge (**Fig. 3C**). As expected, there were little differences in collagen deposition between vehicle-treated *Lrp6*^AKO^ and *Lrp6*^fl/fl^ mice (**Fig. 3A-C**).

**Figure 3.**
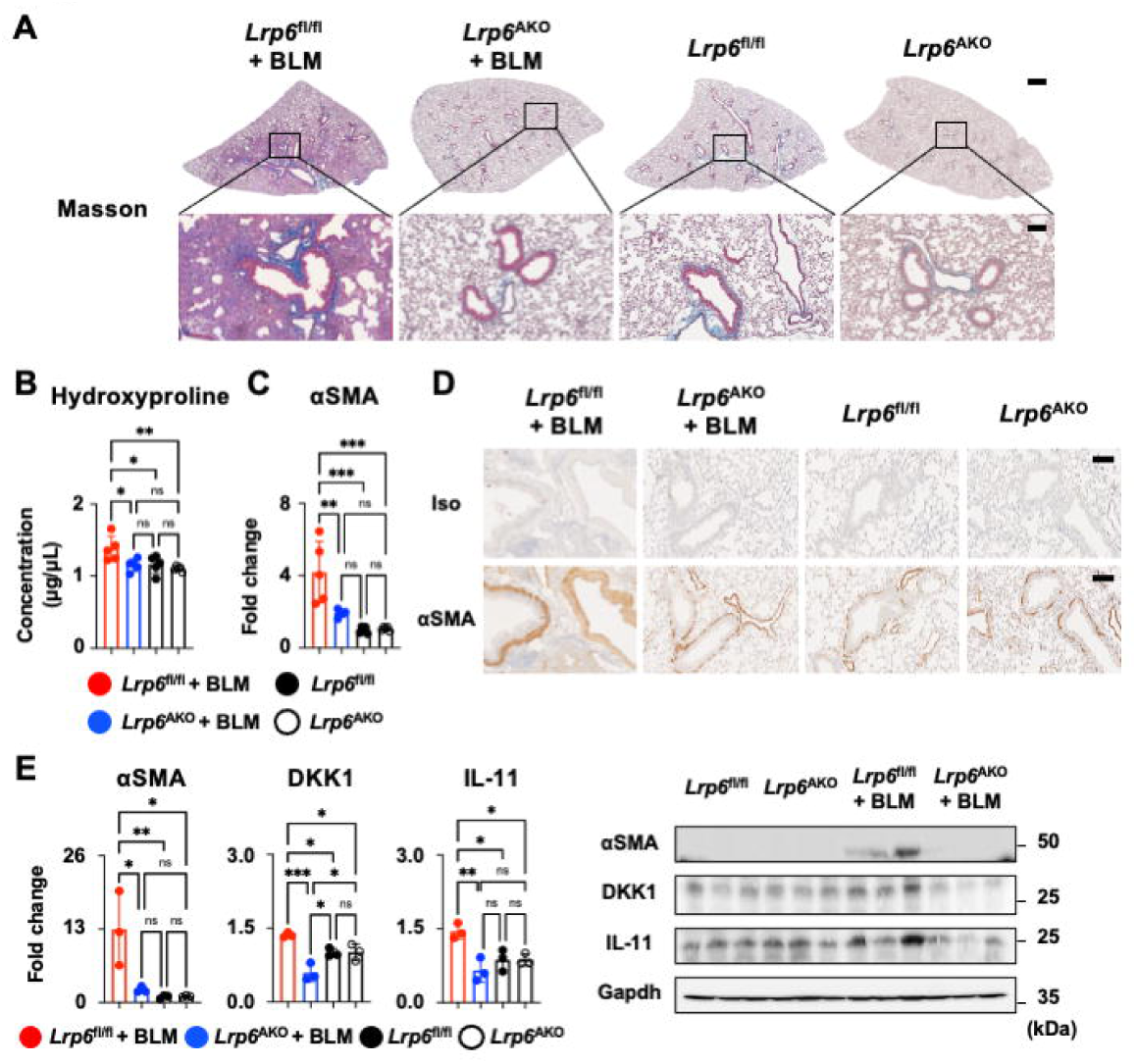
*Lrp6*^AKO^ mice showed reduced BLM-induced lung fibrosis phenotypes. (**A-E**) *Lrp6*^AKO^ and their littermate controls *Lrp6*^fl/fl^ mice were challenged with BLM on day 0. Two weeks later, the lungs were harvested. (**A**) Representative images of Masson staining are shown. Scale bar = 800 μm (top), 100 μm (bottom). (**B**) Hydroxyproline concentration was measured (n = 5). (**C**) αSMA protein levels were analyzed by IHC and ImageJ (n = 5). (**D**) Representative images of IHC are shown. Scale bar = 100 μm. (**E**) The ratio of αSMA, DKK1 and IL-11 to Gapdh was quantified by Western blot and Image J (n = 3). Data are represented as means ± SD. One-way ANOVA with Tukey’s post hoc test. **p < 0.005, *p < 0.05, ns, not significant.

Next, we assessed αSMA protein levels, a myofibroblast marker, in *Lrp6*^AKO^ and *Lrp6*^fl/fl^ mice after the BLM challenge. IHC images showed elevated αSMA protein levels in *Lrp6*^fl/fl^ mice after the BLM challenge, while *Lrp6*^AKO^ mice exhibited a significant decrease (**Fig. 3C, D**). Consistent with these results, Western blot analysis showed a substantial reduction in αSMA protein levels in the lungs of BLM-treated *Lrp6*^AKO^ mice compared to BLM-treated *Lrp6*^fl/fl^ mice (**Fig. 3E**). Vehicle-treated *Lrp6*^AKO^ and *Lrp6*^fl/fl^ mice showed minimal differences in αSMA protein levels (**Fig. 3C-E**).

DKK1 protein levels in the lungs were quantified using Western blot analysis. Previously, it has been reported that DKK1 expression increased after BLM challenge up to day 14 [14,15]. Interestingly, BLM-treated *Lrp6*^AKO^ mice showed a marked decrease in DKK1 protein levels compared to BLM-treated *Lrp6*^fl/fl^ mice, suggesting that the sustained DKK1 expression required LRP6 expression from myofibroblasts (**Fig. 3E**). Given that DKK1 induced *Il11* gene expression in lung fibroblasts in Fig. 1, we measured IL-11 protein levels in *Lrp6*^AKO^ and *Lrp6*^fl/fl^ mice after the BLM challenge. *Lrp6*^AKO^ mice showed a substantial decrease in IL-11 protein levels compared to *Lrp6*^fl/fl^ mice following BLM-induced lung injury (**Fig. 3E**).

Our results suggest that the deletion of LRP6 in fibroblasts abrogates key fibrotic cellular events, such as collagen deposition and myofibroblast activation, thereby protecting mice from lung fibrosis phenotypes upon BLM-induced acute lung injury.

### *Lrp6*^AKO^ mice exhibited markedly reduced lung inflammation upon BLM-induced lung injury

Given that BLM-induced collagen deposition was significantly reduced in *Lrp6*^AKO^ mice, we investigated whether the deletion of LRP6 in αSMA-expressing cells also reduced lung inflammation following the BLM challenge. Two weeks after the BLM challenge, lungs from *Lrp6*^AKO^ and *Lrp6*^fl/fl^ mice were analyzed for histological and immunological phenotypes. H&E staining showed increased inflammation and leukocyte infiltration in the lungs of BLM-treated *Lrp6*^fl/fl^ mice (**Fig. 4A**). BLM-treated *Lrp6*^AKO^ mice exhibited a marked reduction in inflammation and leukocyte infiltration in the lung compared to BLM-treated *Lrp6*^fl/fl^ mice (**Fig. 4A**).

**Figure 4.**
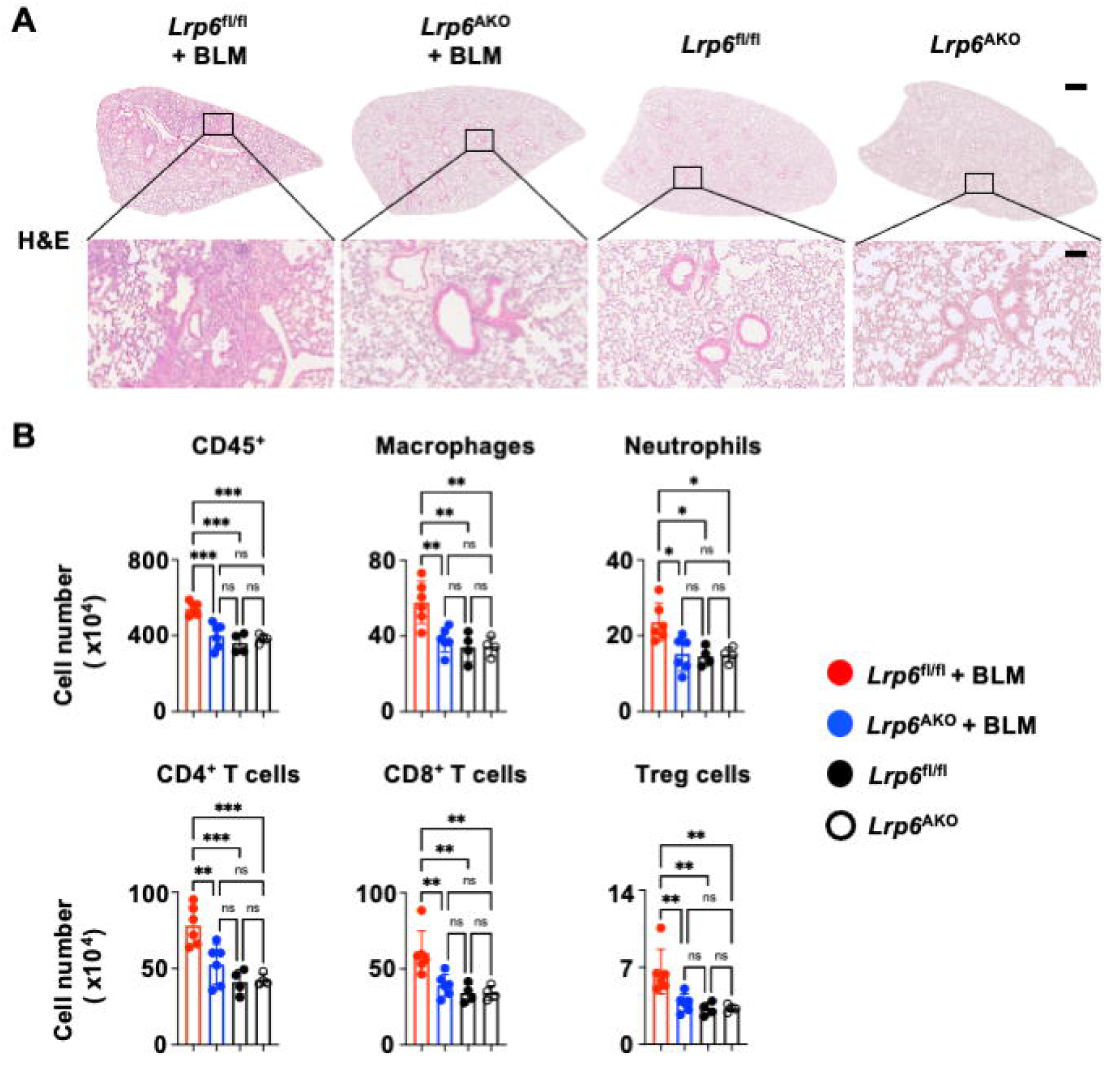
*Lrp6*^AKO^ mice showed reduced BLM-induced lung inflammation. (**A-B**) *Lrp6*^AKO^ and their littermate controls *Lrp6*^fl/fl^ mice were challenged with BLM. Two weeks later, the lungs were harvested. (**A**) Representative images of H&E staining are shown. Scale bar = 800 μm (top), 100 μm (bottom). (**B**) CD45^+^ leukocytes, macrophages, neutrophils, CD4^+^ T cells, CD8^+^ T cells and Treg cells were quantitated by flow cytometry. Data are represented as means ± SD (n = 4 or 6). One-way ANOVA with Tukey’s post hoc test. ***p < 0.001, **p < 0.005, *p < 0.05, ns, not significant.

Next, immune cell populations from lung tissue homogenates were quantitated by flow cytometry (**Fig. S3**). Consistent with the H&E staining results, leukocyte infiltration was substantially decreased in *Lrp6*^AKO^ mice compared to *Lrp6*^fl/fl^ mice upon BLM challenge (**Fig. 4B**). BLM-treated *Lrp6*^fl/fl^ mice showed increased numbers of macrophages, neutrophils, CD4^+^ T cells, CD8^+^ T cells and Treg cells (**Fig. 4B**). These increases in immune cell population upon BLM injury were significantly abolished in *Lrp6*^AKO^ mice compared to *Lrp6*^fl/fl^ mice in the lung (**Fig. 4B**). No notable differences were observed in immune cell infiltration and populations between vehicle-treated *Lrp6*^AKO^ and *Lrp6*^fl/fl^ mice (**Fig. 4A, B**). Our results demonstrated that the ablation of LRP6 in αSMA-expressing cells abrogated lung inflammation upon BLM challenge.

## Discussion

In this study, we showed that genetic deletion of LRP6 in αSMA-expressing cells using *Lrp6*^AKO^ mice abrogated lung inflammation and fibrosis phenotypes in a BLM-induced lung injury mouse model. The ablation of LRP6 in αSMA-expressing cells showed very little effect on baseline immune cell profiles and collagen deposition under homeostatic conditions. DKK1 induced genes important for inflammation and tissue fibrosis in lung fibroblasts, highlighting the role of LRP6 in αSMA-expressing cells in regulating lung inflammation and fibrosis in the BLM model.

Previously, our studies showed that *Dkk1* hypomorphic *doubleridge* mice and thrombocyte-specific *Dkk1* deletion protected from BLM-induced lung injury, indicating that prolonged DKK1 expression following lung injury promotes inflammation and fibrosis [14,15]. Our findings showed that DKK1 induced gene profiles related to inflammation and fibrosis in lung fibroblasts and *Lrp6*^AKO^ mice had reduced lung inflammation and lower DKK1 protein levels following the BLM challenge. These results suggest that the ablation of LRP6 in lung fibroblasts inhibited prolonged DKK1 expression and its pro-inflammatory role in BLM lung injury. Further studies are warranted to explore the mechanisms through which DKK1 modulates fibroblasts to promote lung inflammation and fibrosis.

It has been shown that gene expression of *IL11* is significantly upregulated in human PF lungs [35]. Both systemic recombinant IL-11 administration and fibroblast-specific *Il11* over expression were sufficient to drive lung fibrosis phenotypes in mice, indicating that IL-11 is an important driver of lung fibrosis [35]. Our results showed that DKK1 induced *Il11* gene expression in lung fibroblasts and BLM-treated *Lrp6*^AKO^ mice exhibited decreased IL-11 protein expression levels, highlighting the role of the DKK1-LRP6 axis in modulating IL-11 expression. Further studies are needed to investigate how DKK1 regulates IL-11 expression via LRP6 in fibroblasts. Given that overexpression of *Il11* in smooth muscle cells can cause cross-tissue inflammation and fibrosis [36], more studies are warranted to understand the role of the DKK1-LRP6-IL-11 axis in different types of injuries and organs.

Our previous research showed that the deletion of LRP6 in the myeloid cell lineage reduced inflammation, collagen deposition and αSMA protein levels in BLM-induced lung injury [50]. Consistently, BLM-treated *Lrp6*^AKO^ mice exhibited similar results, suggesting the importance of the DKK1-LRP6 axis in regulating crosstalk between macrophages and fibroblasts during BLM-induced lung injury. Given that thrombocyte-derived DKK1 modulated macrophages in BLM-induced lung injury [15], it would be worth to investigate whether thrombocytes communicate with macrophages and fibroblasts via DKK1 to orchestrate inflammation-induced injury repair process.

Transforming growth factor β1 (TGFβ1) is a profibrotic cytokine that promotes PF by inducing the differentiation and activation of αSMA^+^ myofibroblast [51–54]. It has been shown that reduced DKK1 expression in *doubleridge* mice led to decreased TGFβ1 protein levels upon BLM challenge in the lung [14]. Our results showed that DKK1 induced gene expressions known to promote inflammation and fibrosis in lung fibroblasts. Further studies investigating the interplay between DKK1 and TGFβ1 for lung fibroblasts are warranted to provide more insights and new strategies for treating lung fibrosis.

Our study has limitations in that αSMA expression is not exclusively limited to myofibroblasts although αSMA is used as a marker for activated myofibroblasts. Further studies using other fibroblast-specific Cre strains such as *Fsp1*-cre or *Postn*-cre are warranted to gain more comprehensive insights into the role of LRP6 in lung fibroblasts.

Taken together, our study showed that LRP6 in lung fibroblasts expresses proinflammatory and profibrotic genes, and it plays a crucial role in regulating lung inflammation and fibrosis upon BLM-induced lung challenge. This places the DKK1-LRP6 axis as a key modulator of inflammation and fibrosis, emphasizing its potential as a therapeutic target for lung fibrosis.

## Abbreviations

LRP6: Low-density lipoprotein receptor-related protein 6
DKK1: Dickkopf1
BLM: Bleomycin
PF: Pulmonary fibrosis
αSMA: Alpha-smooth muscle actin
ECM: Extracellular matrix

## Acknowledgements

The author(s) declare financial support was received for the research, authorship, and/or publication of this article. This work was supported in part by the Massey Comprehensive Cancer Center Innovation Pilot Award (2023-INN-CB, to W-J.C.), and the Virginia Commonwealth University fund (W-J.C.), VCU’s CTSA UL1TR002649 from the National Center for Advancing Translational Science and the CCTR Endowment Fund of Virginia Commonwealth University (W-J.C.), and in part by Institutional Research Grant (IRG-18-159-43 and IRG-21-134-46) from the American Cancer Society (W-J.C.). Histology Services in support of the research project were generated by the Virginia Commonwealth University Cancer Mouse Models Core Laboratory, supported, in part, with funding from NIH-NCI Cancer Center Support Grant P30 CA016059.

## Author contributions

E-A.S., S. S. and W-J.C. designed experiments, analyzed data, and wrote the manuscript. E-A.S., S. S., T. A., and M.H.P performed experiments. M. G. D. analyzed RNA-seq data. P. J. S. gave input and guidance on the lung injury model. W-J.C. supervised all aspects of the project. All of the authors read and commented on the manuscript.

## Data accessibility

The data that support the findings of this study are available in the main text, figures, and the supporting information of this article. The raw bulk RNA seq data presented in this study is deposited in the NCBI GEO repository (accession number GSE276111, https://www.ncbi.nlm.nih.gov/geo/).

## Notes

### Competing Interest Statement

The authors have declared no competing interest.

