## Supplementary Figures for "Ablation of LRP6 in alpha-smooth muscle actin-expressing cells abrogates lung inflammation and fibrosis upon bleomycin-induced lung injury"

**Figure S1.**

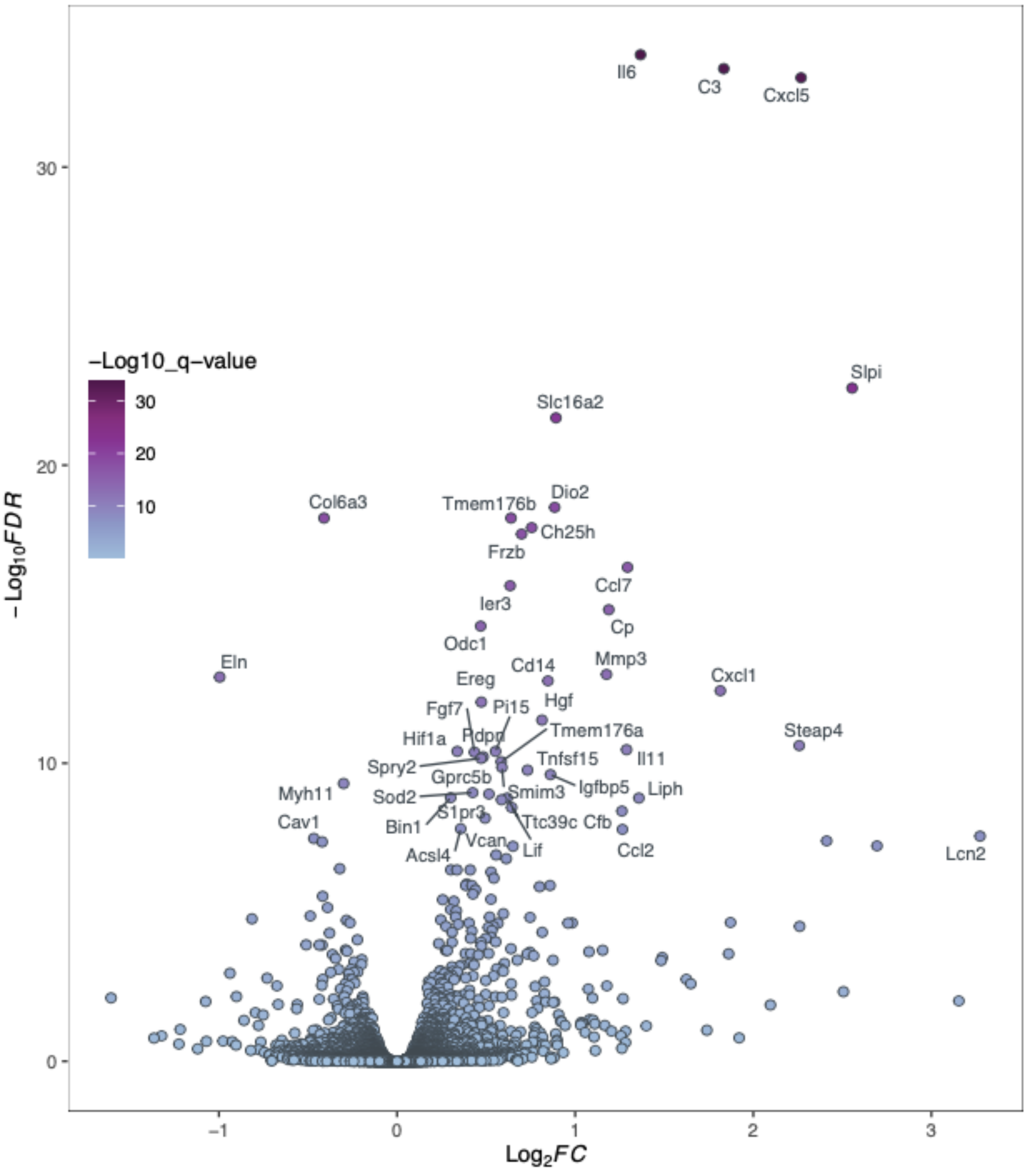


**Supplementary Fig. 1:** Lung fibroblasts were treated with DKK1 for 24 hours prior to bulk RNA seq. DEG analysis between untreated and DKK1-treated group is visualized in the form of a volcano plot.

**Figure S2.**

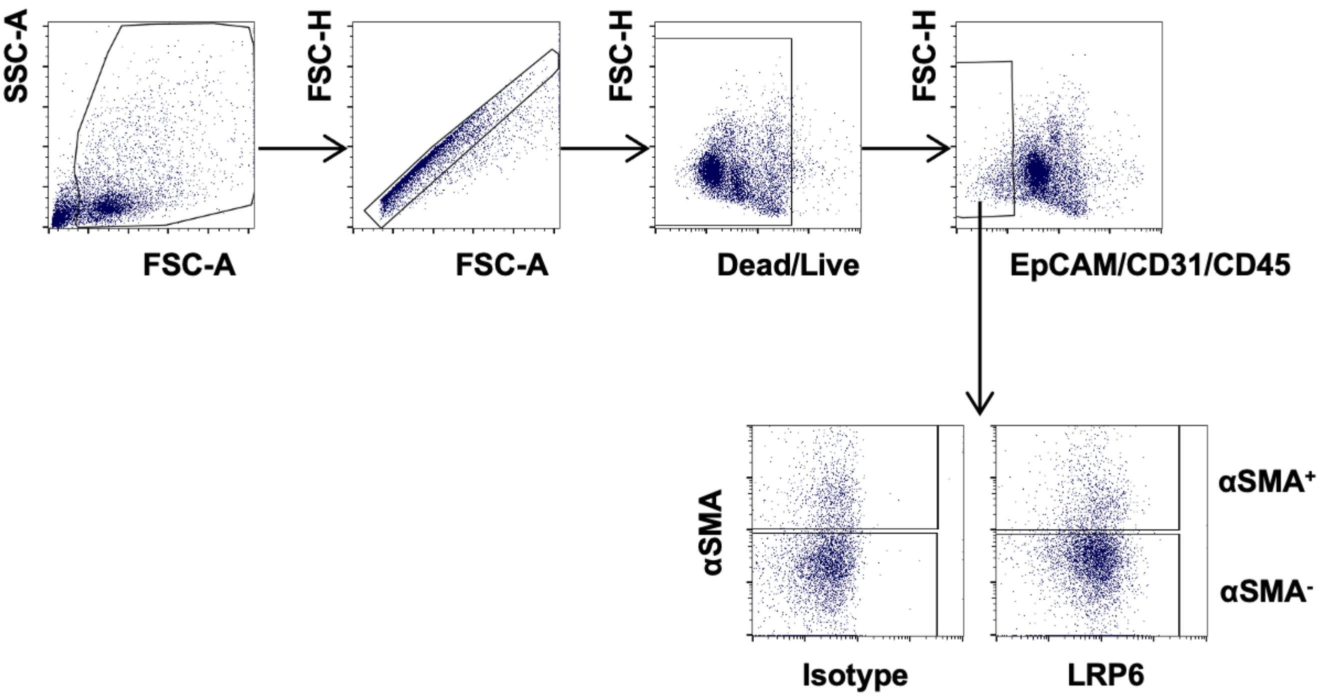


**Supplementary Fig. 2:** Flow cytometry gating strategy used for identification of LRP6 expressions in αSMA$^{+}$ or αSMA$^{-}$ populations in mouse lung homogenates is shown.

**Figure S3.**

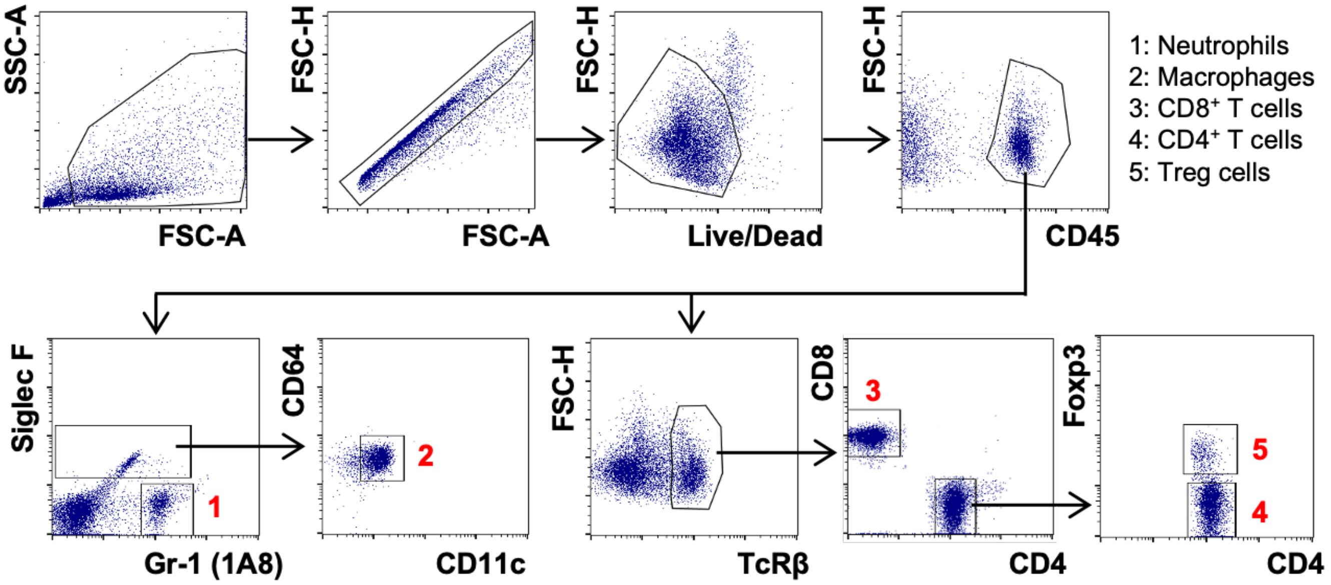


**Supplementary Fig. 3:** Flow cytometry gating strategy used for immune profiles in mouse lung homogenates is shown.
