## Supplemental Table S2 for "Ablation of LRP6 in alpha-smooth muscle actin-expressing cells abrogates lung inflammation and fibrosis upon bleomycin-induced lung injury"

Information on Antibodies

| Antibody | Manufacturer | Catalog No. |
|---|---|---|
| CD45 Monoclonal Antibody (30-F11), Biotin | eBioscience | 13-0451-82 |
| Biotin anti-mouse CD31 Antibody | BioLegend | 102503 |
| Biotin anti-mouse CD326 (Ep-CAM) Antibody | BioLegend | 118203 |
| Biotin anti-mouse CD206 (MMR) | BioLegend | 141714 |
| eFluor™ 450 Streptavidin | eBioscience | 48-4317-82 |
| Pacific Blue™ anti-mouse TCR beta chain, clone H57-597 | BioLegend | 109226 |
| Pacific Blue™ Rat IgG2α kappa Isotype Ctrl Antibody, clone RTK2758 | BioLegend | 400527 |
| FITC anti-mouse Ly-6G | BioLegend | 127605 |
| CD4 Monoclonal Antibody (RM4-5), Biotin | eBioscience | 13-0042-82 |
| FITC Streptavidin | BioLegend | 405202 |
| FITC Mouse IgG1 kappa Isotype Ctrl Antibody, clone MOPC-21 | BioLegend | 400107 |
| Collagen Type I Antibody FITC Conjugated | ROCKLAND | 600-402-103 |
| PE anti-mouse TLR4 (CD284)/MD2 Complex | Biolegend | 117605 |
| Arginase 1 Monoclonal Antibody (A1exF5), PE | eBioscience | 12-3697-80 |
| FOXP3 Monoclonal Antibody (FJK-16s), eFluor™ 660 | eBioscience | 50-5773-80 |
| PerCP/Cyanine5.5 anti-mouse CD64 (FcγRI) | BioLegend | 139308 |
| PerCP/Cyanine5.5 anti-mouse CD8b.2 | BioLegend | 140417 |
| PerCP/Cyanine5.5 Rat IgG2a, κ Isotype Ctrl | BioLegend | 400531 |
| PE/Cyanine7 anti-mouse CD11c | BioLegend | 117317 |
| NK1.1 Monoclonal Antibody (PK136), PE-Cyanine7 | eBioscience | 25-5941-82 |
| PE/Cyanine7 anti-mouse CD140a | BioLegend | 135911 |
| Ki-67 Monoclonal Antibody (SolA15), PE-Cyanine7 | Invitrogen | 25-5698-82 |
| PE/Cyanine7 Mouse IgG1, κ Isotype Ctrl | BioLegend | 400125 |
| Arginase 1 Monoclonal Antibody (A1exF5), APC | Invitrogen | 17-3697-82 |
| APC anti-mouse CD284 (TLR4) | Biolegend | 145405 |
| Alpha-Smooth Muscle Actin Monoclonal Antibody (1A4), eFluor™ 660 | eBioscience | 50-9760-80 |
| APC Mouse IgG1 κ Isotype Control | BD Biosciences | 550854 |
| APC/Fire™ 750 anti-mouse CD45, clone 30-F11 | Biolegend | 103154 |
| CD170 (Siglec F) Monoclonal Antibody (1RNM44N), Super Bright™ 600 | eBioscience | 63-1702-82 |
| Rat IgG2a kappa Isotype Control (eBR2a), Super Bright™ 600 | eBioscience | 63-4321-80 |
| Ultra-LEAF™ Purified anti-mouse CD16/32 Antibody | BioLegend | 101330 |
